# Expanding adult tubular microvessels on stiff substrates with endothelial cells and pericytes from the same tissue

**DOI:** 10.1101/2022.04.12.488033

**Authors:** Xiuyue Song, Yali Yu, Jie Mu, Zihan Wang, Yu Leng, Yalan Xu, Hai Zhu, Xuefeng Qiu, Peifeng Li, Jing Li, Dong Wang

## Abstract

Microvessels are essential for tissue engineering and regeneration. In current methods, endothelial cells are usually cultured in commercially available media and form a monolayer of cell sheets on stiff substrates and a tubular structure when cultured with soft hydrogels. To mimic the microvessels *in vivo*, researchers usually coculture the endothelial cells and pericytes from different adult tissues or derived from pluripotent stem cells in a three-dimensional hydrogel. However, there is a challenge for these models to reflect tissue-specific characteristics due to the vascular heterogeneity throughout the body. Here, we established a culture model for expanding adult tubular microvessels on stiff substrates with endothelial cells and pericytes derived from the same tissue. We isolated microvessels from adult rat subcutaneous soft connective tissue and cultured them on regular plastic dishes. We performed a series of screenings and formulated a custom-made medium (Medium-X), containing mainly antioxidants and three small molecules, Chir99021, A83-01, and Y27632. Medium-X significantly promoted adult microvessel growth while maintaining their characteristic tubular morphology up to 8 weeks *in vitro*, contrary to the monolayer of endothelial cell sheets in the commercially available medium EGM2MV. Transcriptomic analysis showed that Medium-X maintained the tubular morphology of microvessels by promoting angiogenesis and vascular remodeling while suppressing oxidation and lipid metabolic pathways. The model presented in this study can be applied to other organs for expanding organ-specific microvessels for tissue engineering and vascular regeneration.

## 1. Introduction

Microvessels play essential roles in maintaining normal organ functions. Microvascular dysregulation happens in various diseases, such as fibrosis, tumor, and diabetic microvasculopathy^1^. Microvascular structural stability and proper function rely on microvascular endothelial cells (ECs) and pericytes^2^. Although many studies have focused on embryonic microvascular development using animal models, *in vitro* culture models of adult microvessels will provide more insights into the mechanisms of microvascular diseases in adulthood. The isolation and growth of microvessels from adult tissues can provide a platform for research on patient-specific diseases and benefit the development of precision medicine.

There have been various *in vitro* culture models for adult microvessels^3^. Active angiogenesis can be induced in tissue explant culture models, such as retina and aorta ring explants in a three-dimensional (3D) matrix^3^; however, these models face challenges while being translated into clinical settings. It has been widely used to culture a pure population of primary ECs in most laboratories. The human umbilical vein is a popular source for primary EC isolation. ECs can also be isolated from other organs, e.g., the kidney, then purified by flow cytometric sorting^4,5^. The co-culture models of ECs and pericytes have been developed to investigate the mechanisms of pericyte recruitment. The pericytes used in these experiments were usually isolated from the brain, an organ rich in pericytes^6^. In recent years, pluripotent stem cells (PSCs) have been an emerging source for producing ECs, pericytes, and smooth muscle cells^7^. However, it is a challenge for the vascular cells derived from PSCs to reflect the biological characteristics of a specific tissue or organ. Due to the heterogeneity of blood vessels throughout the body^8^, the ideal culture models for adult microvessels would contain microvascular ECs and pericytes from the same adult tissue, which will benefit organ-specific studies.

In traditional methods, ECs are usually cultured in commercially available media and form a monolayer on the rigid surface of plastic dishes^3^. Many efforts have been made to achieve a tubular morphology of microvessels *in vitro* to recapitulate the microvascular environment *in vivo*. ECs can form a network of tubular microvessels on the surface of a soft hydrogel, e.g., Matrigel, which, however, can only be maintained for a few days^3^. Tubular microvessels can also be obtained by 3D culture of ECs inside hydrogels, such as collagen, fibrin, Matrigel, and synthetic hydrogels^3,9^. In recent years, micro-technologies, such as microfluidics, 3D printing, and ice templating^9^, make it possible to fabricate elegant microvascular models. ECs can be seeded into the microchannels made by extracellular protein hydrogels and adhered to the channel walls to form a tubular vessel^10^. ECs embedded in 3D hydrogels can self-assemble to form microvessels, further induced to form a perfused microvascular network in microfluidic channels^11^. These microvessels in 3D hydrogels were usually maintained *in vitro* for 2 weeks. Although superior to 2D culture in forming tubular microvessels, 3D cultures face difficulties in characterizing the microvessels by conventional biochemical methods, such as immunostaining and confocal imaging. In addition, hydrogels are expensive for most laboratories.

In this proof-of-concept study, we aimed to establish a culture model for expanding adult tubular microvessels with syngeneic pericytes on the rigid surface of regular plastic dishes. Microvessels were isolated from adult rats’ subcutaneous soft connective tissue and cultured in a custom-made microvessel medium. By activating Wnt and inhibiting TGFβ and ROCK signaling pathways, we expanded tubular microvessels wrapped with syngeneic pericytes on regular plastic culture dishes. The tubular microvessels can be maintained for up to 8 weeks *in vitro*.

## 2. Results

### 2.1 Expanding tubular adult microvessels in Medium-X

Primary microvessels were isolated from adult Sprague Dawley (SD) rats (Fig. 1, A). The subcutaneous soft connective tissue was harvested from adult rats and digested in an enzymatic solution containing mainly collagenase I at 37 °C. We designed a cyclic digestion method to improve the viability of primary microvessels (Fig. 1, A). Every 10 minutes, we harvested cell suspension from the digestion tube and added a freshly prepared enzymatic solution to the tube. The cell suspension was centrifuged, and the pellets were re-suspended in a collection tube containing phosphate-buffered saline (PBS). The digestion cycle continued until all the tissue pieces were wholly digested. The cell suspensions were pooled together and filtered through a 30 μm strainer. The microvessel segments were collected from the strainer mesh (Fig. 1, A). We obtained the microvessel segments with a length range of about 50 - 150 μm (Fig. 1, B; Supplemental Fig. S1). In the commercially available medium, EGM2MV, ECs formed a monolayer on the rigid surface of plastic culture dishes, consistent with traditional culture methods.

**Fig 1.**
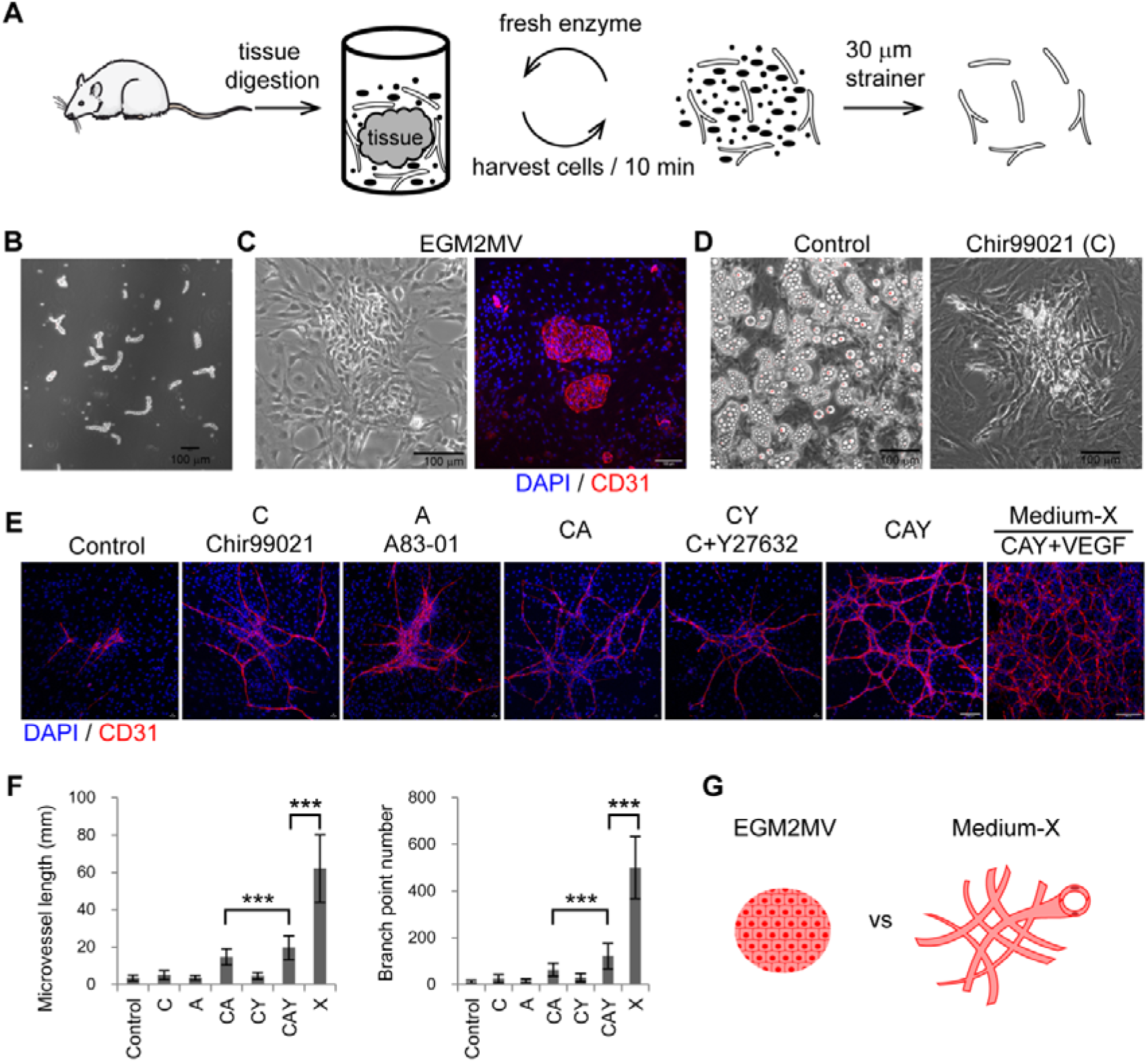
Expand adult tubular microvessels *in vitro*. A, The procedure of isolating microvessels from adult SD rats’ subcutaneous soft connective tissues by cyclic enzymatic digestion. B, Phase contrast image of primary microvessels. C, Phase contrast and immunofluorescence images of the microvessels cultured in EGM2MV medium. D, Phase-contrast images of the primary microvessels cultured in the media of control and Chir99021. E, Immunofluorescence images of primary microvessels cultured in different media. F, Quantification of the microvessels in different experimental groups including control (n=81), C (n=84), A (n=72), CA (n=97), CY (n=58), CAY (n=61), and Medium-X (CAY+VEGF) (n=9). Data were presented as mean ± SD. One-way ANOVA was performed on the data, followed by Bonferroni post hoc tests. ***, p < 0.001. G, Illustration of the morphologies of microvessels cultured in EGM2MV and Medium-X. The antibody against CD31 was used to label endothelial cells. DAPI was used to label cell nuclei. Scale bars, 100 μm.

To make a better culture system, we set up to formulate a basal medium that was DMEM/F12 supplemented with some ingredients. Considering that oxidative stress was the primary threat to the cells *in vitro*, we tested the effect of adding several antioxidants, such as nicotinamide, NAC, glutathione, and Vitamins C, to the DMEM/F12 medium (See Materials and methods). We then screened signaling pathways that may impact the growth of adult microvessels *in vitro*. Vascular endothelial growth factor (VEGF) was not added in this screening because its potent proangiogenic effect may mask any roles of other signaling pathways. During 2 weeks of culture *in vitro*, most groups’ culture dishes were covered by lipid-like cells characterized by the accumulation of lipid-like droplets, except those supplemented with Chir99021 (C) (Fig. 1, D). Chir99021 is a GSK3β inhibitor and thus stabilizes β-catenin and activates the canonical Wnt signaling pathway. This result was consistent with a previous report that upregulated Wnt signaling inhibited adipogenesis^12^. Thus, Chir99021 was added to the basal medium for the following study.

ECs may undergo endothelial-to-mesenchymal transition and lose EC identity, during which the TGFβ signaling pathway is a master regulator^13^. We thus examined the effect of an inhibitor of the TGFβ signaling pathway, A83-01 (A), in combination with Chir99021. We further examined another small molecule, Y27632 (Y), a ROCK inhibitor that was reported to improve embryonic stem cell survival *in vitro*^14^. The different combinations of these three small molecules were investigated (Fig. 1, E). The primary microvessels were cultured for about a week and analyzed by immunofluorescence staining. We found that the combination of the three small molecules, C+A+Y, significantly promoted the growth of tubular microvessels (Fig. 1, E and F). The addition of VEGF further significantly increased microvessel length and branch point number (Fig. 1, E and F). We referred to the medium containing C, A, Y, and VEGF as Medium-X, which significantly promoted the growth of adult tubular microvessels *in vitro*, compared to the commercially available medium EGM2MV (Fig. 1, G).

### 2.2 Medium-X upregulated angiogenesis and downregulated oxidation and lipid metabolism

We performed mRNA sequencing to analyze gene expression profiles of the microvessels in Medium-X (X) and EGM2MV (E) compared to fresh primary microvessels (V). We focused on the genes up- or down-regulated in EGM2MV and Medium-X (Supplemental Fig. S2, A). KEGG pathway analysis showed that EGM2MV specifically upregulated the pathways of oxidation and lipid metabolism (Supplemental Fig. S2, B). GO analysis showed that the elevated biological processes in EGM2MV included mainly oxidation, lipid metabolism, and aging, which involved increased mitochondrial activities (Fig. 2, A; Supplemental Fig. S2, C). On the contrary, oxidation- and lipid metabolism-related processes were significantly suppressed in Medium-X (Supplemental Fig. S2, A). Medium-X specifically upregulated Notch, TNF, Rap1, Ras, PI3K-Akt, and chemokine signaling pathways, which were closely related to angiogenesis (Supplemental Fig. S2, B). GO analysis showed that the upregulated biological processes in Medium-X included mainly angiogenesis, vascular branch formation, vascular remodeling, EC proliferation, and migration, which mainly involved cell-cell junctions between ECs, contributing to the structural integrity of microvessels (Fig. 2, B; Supplemental Fig. S2, D).

**Fig. 2.**
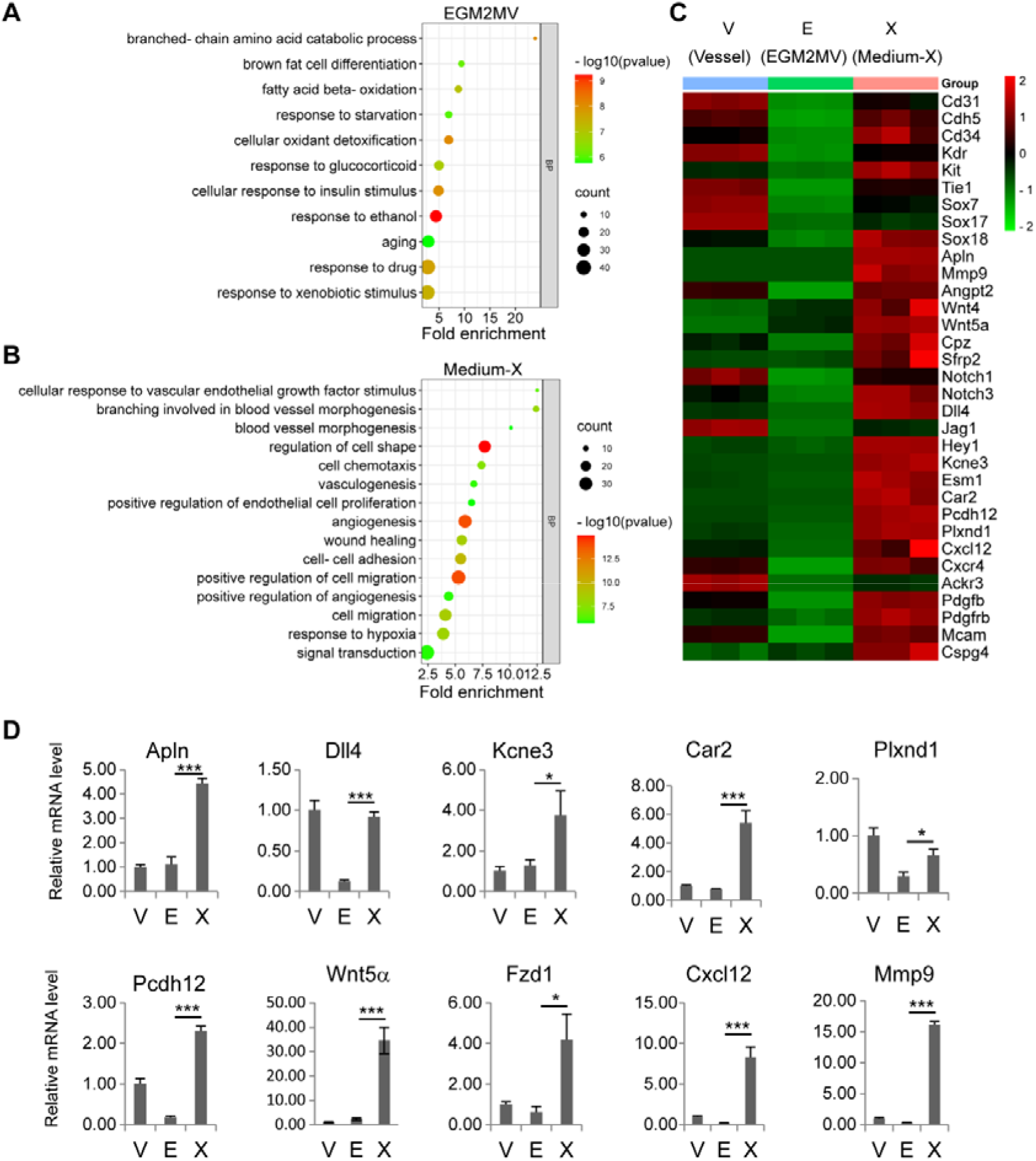
Transcriptomic analysis of Medium-X. A and B, GO analysis of the biological processes upregulated in EGM2MV (A) and Medium-X (B). C, The heatmap of the genes that were downregulated in EGM2MV and upregulated in Medium-X. D, Quantitative RT-PCR verification of gene expression in different groups. Data were presented as mean ± SD. One-way ANOVA was performed on the data, followed by Bonferroni post hoc tests. *, p < 0.05, ***, p < 0.001.

The genes for the identity of ECs (Cd31, Cdh5, Tie1^15,16^) and endothelial progenitor cells (Cd34, Kdr, and Kit)^17^ were down-regulated in EGM2MV but significantly upregulated in Medium-X (Fig. 2, C). Sox family members, including Sox7, 17, and 18, were also upregulated in Medium-X for the differentiation and maturation of ECs^18^. Medium-X also upregulated the expression of Apln, which was recently reported to mark an EC subpopulation contributing to vascular development^19,20^ and regeneration^21,22^. Proangiogenic genes were significantly upregulated in Medium-X, such as Mmp9^23^, Angpt2^24^, Wnt signaling genes (Wnt4, Wnt5a, Cpz, Sfrp2)^25^, and Notch signaling genes (Notch1, Notch3, Dll4, Jag1, Hey1)^26–28^, and chemokine signaling (Cxcl12, Cxcr4, and Ackr3)^29,30^ (Fig. 2, C and D). Genes significantly upregulated in Medium-X also included: Kcne3^31^, Esm1^32^, as tip EC markers; Car2, which promoted EC survival^33^; Pcdh12, a protocadherin^34^; Plxnd1, for EC mechanotransduction^35^.

In addition to the markers of ECs, Medium-X also upregulated Pdgfb and Pdgfrb, which were expressed in ECs and pericytes respectively^36^. Other pericyte markers, including Mcam (Cd146) and Cspg4 (Ng2)^37^ were also upregulated in Medium-X. These results indicated that both ECs and pericytes of the same microvessel segments were expanded in Medium-X.

### 2.3 Individual roles of Chir99021, Y27632, and A83-01 in tubular microvessel growth in vitro

To investigate the roles of the small molecules in Medium-X, VEGF was supplemented to the basal medium to obtain enough microvessels for analysis. The primary microvessels were cultured in different groups of the combinations of the three small molecules. During a week of culture *in vitro*, a single small molecule Chir99021 alone was enough to promote tubular microvessels’ growth significantly. Adding Y27632 further increased microvessel length and branch point number. Adding A83-01 to C or CY groups did not benefit microvessel length or branch point number but produced some local areas with wide EC bundles (Fig. 3, A and B).

**Fig. 3.**
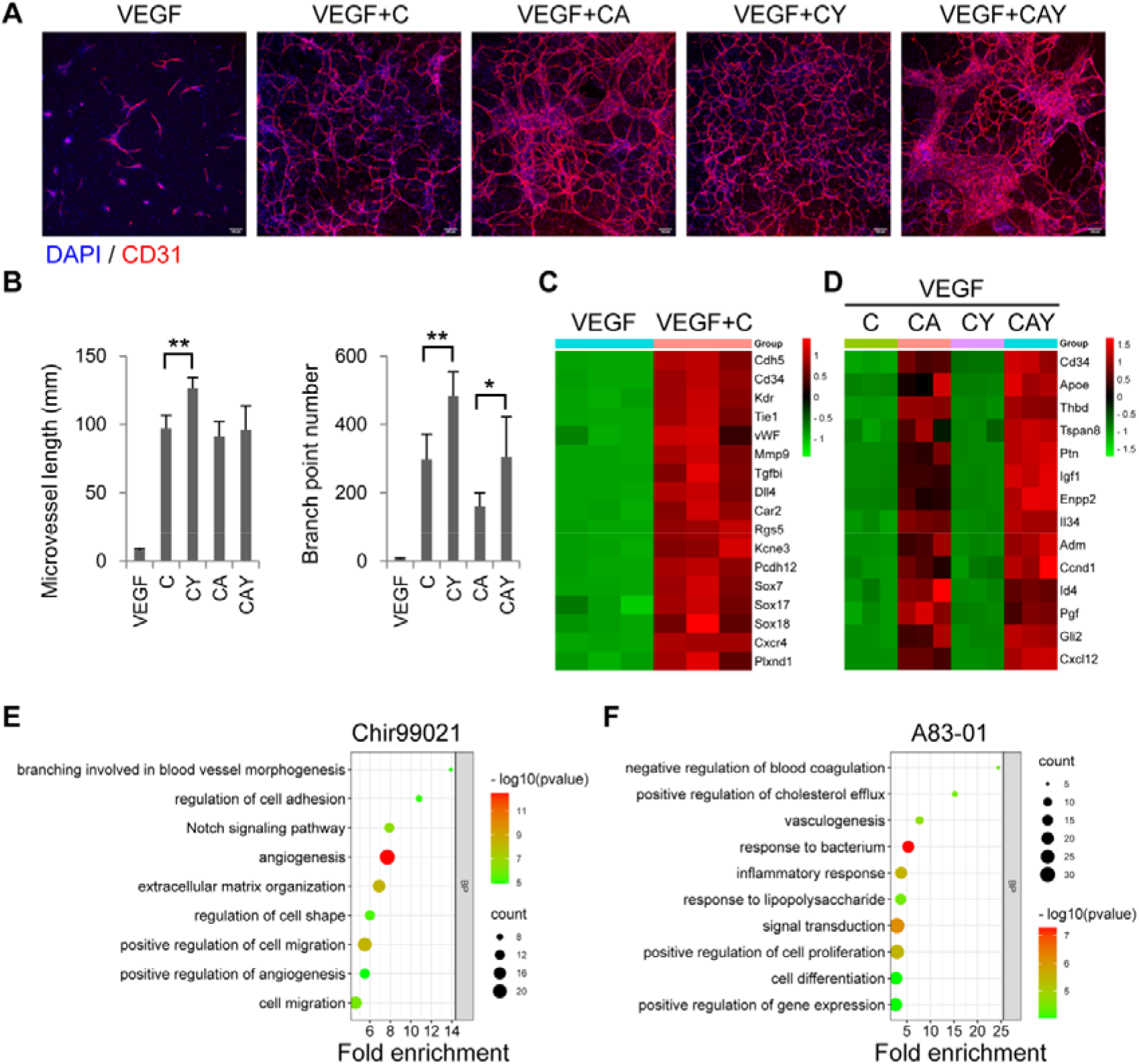
The roles of individual small molecules. A, Immunofluorescence images of the microvessels in different groups. The antibody against CD31 was used to label endothelial cells. DAPI was used to label cell nuclei. Scale bars, 100 μm. B, Quantification of microvessel length and branch point number in different groups (n = 6). Data were presented as mean ± SD. One-way ANOVA was performed on the data, followed by Bonferroni post hoc tests. *, p < 0.05, **, p < 0.01. C and D, The heatmaps of genes upregulated by Chir99021 (C) and A83-01 (D) in different groups. E and F, GO analysis of the biological processes upregulated by Chir99021 (E) and A83-01 (F).

Transcriptomic analysis showed that the upregulated expression of most proangiogenic genes was mainly attributed to Chir99021, which acted as a Wnt agonist and induced the activation of the Notch signaling and other pathways associated with angiogenesis, branch formation, and cell migration (Fig. 3, C and E). A83-01 played a minor role in angiogenesis but contributed to preserved EC functions by upregulating the expression of genes related to cholesterol efflux (Apoe), anti-coagulation (Thbd), and cell proliferation (such as Pgf, Igf1, and Adm.) (Fig. 3, D and F). Both Chir99021 and A83-01 promoted the expression of the progenitor marker CD34, indicating that EC progenitor properties were maintained and Endo-MT was suppressed (Fig. 3, D). Interestingly, the addition of Y27632 did not significantly change the gene expression profiles of the microvessels (data not shown), although there was indeed an improvement in the microvascular length and branch formation (Fig. 3, A and B). Y27632 may only take effect at the protein level through regulating cell contractility.

### 2.4 Long-term culture of adult tubular microvessels in vitro

We next examined the long-term performance of the three small molecules in the growth of microvessels *in vitro*. For a total of 8 weeks of culture, the cells were fixed and immunostained for EC marker CD31 at different time points: 1, 2, 3, 4, and 8 weeks. Although Chir99021 significantly promoted the growth of microvessels during the first week (Fig. 3), the microvessels in the Chir99021 medium degraded quickly from the second week (Fig. 4, A, E, and F). The addition of A83-01 to Chir99021 medium slightly rescued the microvessels from the second week (Fig. 4, B, E, and F). Y27632 had a noticeable effect on the survival of microvessels from the second week (Fig. 4, C-F). The full Medium-X, containing C, A, and Y, could sustain microvascular growth for 8 weeks (Fig. 4, D-F). Microvascular length and branch formation were relatively high for at least 4 weeks (Fig. D-F). The wide EC bundles in the groups of CA and CAY disappeared, and all the ECs adopted a tubular structure from the second week, indicating that active microvascular remodeling happened during culture *in vitro* (Fig. 4, B and D).

**Fig 4.**
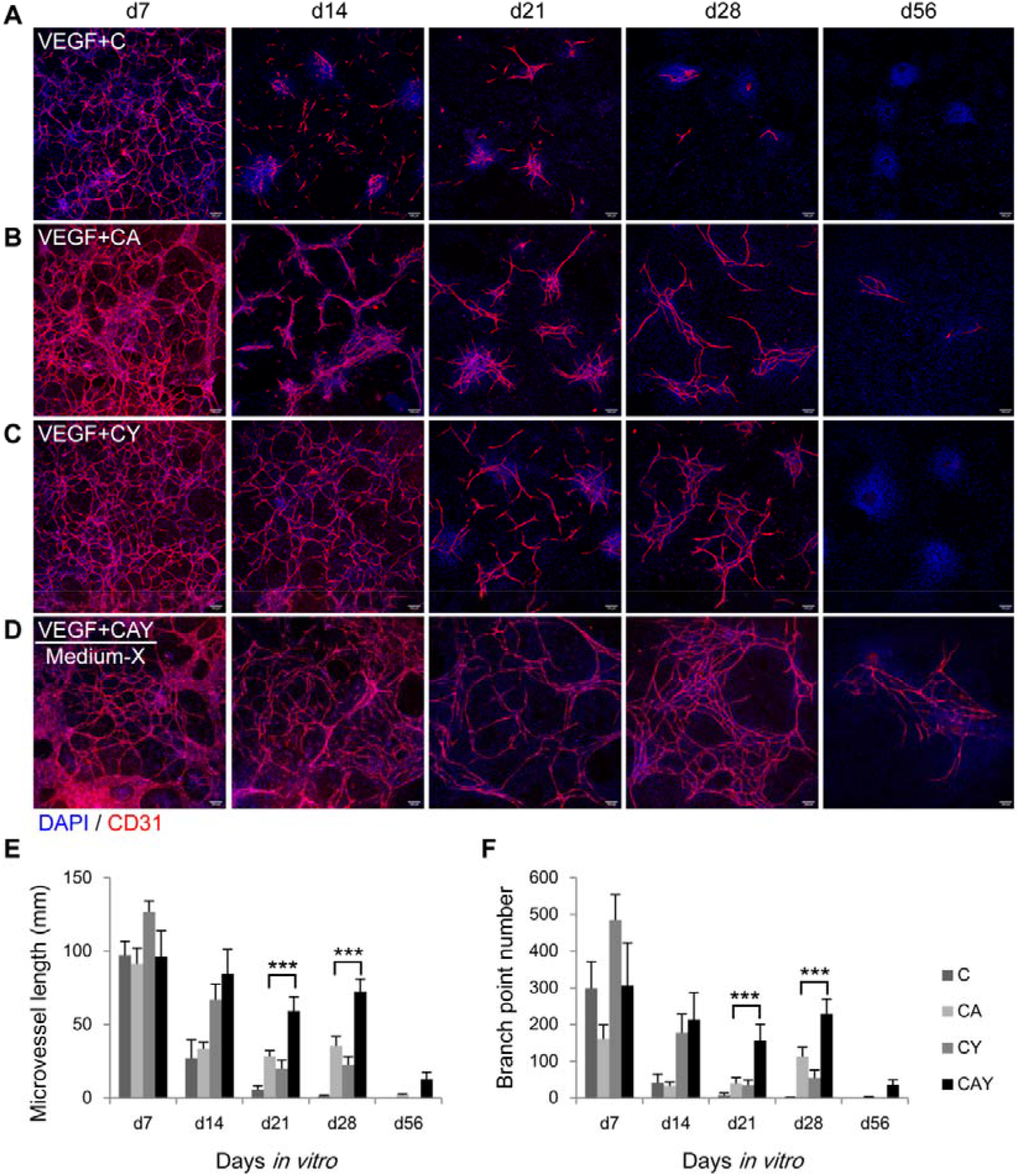
Time-lapse analysis of the microvessels in different media. A-D, Immunofluorescence images of the microvessels in the media supplemented with C (A), CA (B), CY (C), and CAY (D) in the presence of VEGF. The antibody against CD31 was used to label endothelial cells. DAPI was used to label cell nuclei. Scale bars, 100 μm. E and F, Quantification of microvessel length and branch point number in different groups at different time points (n = 6). Data were presented as mean ± SD. Two-way ANOVA was performed on the data, followed by Bonferroni post hoc tests. ***, p < 0.001.

### 2.5 Preserving syngeneic pericytes around tubular microvessels in vitro

There were few LYVE1+ lymphatic endothelial cells in the culture, indicating that our custom-made Medium-X could specifically expand tubular blood vessels (Fig. 5, A). The adult tubular microvessels also showed strong endothelial marker vWF (Fig. 5, B) and the tight junction protein ZO1 (Fig. 5, C). More importantly, plenty of pericytes around the microvessels expressed markers of NG2, PDGFRβ, and αSMA (Fig. 5, D-F).

**Fig 5.**
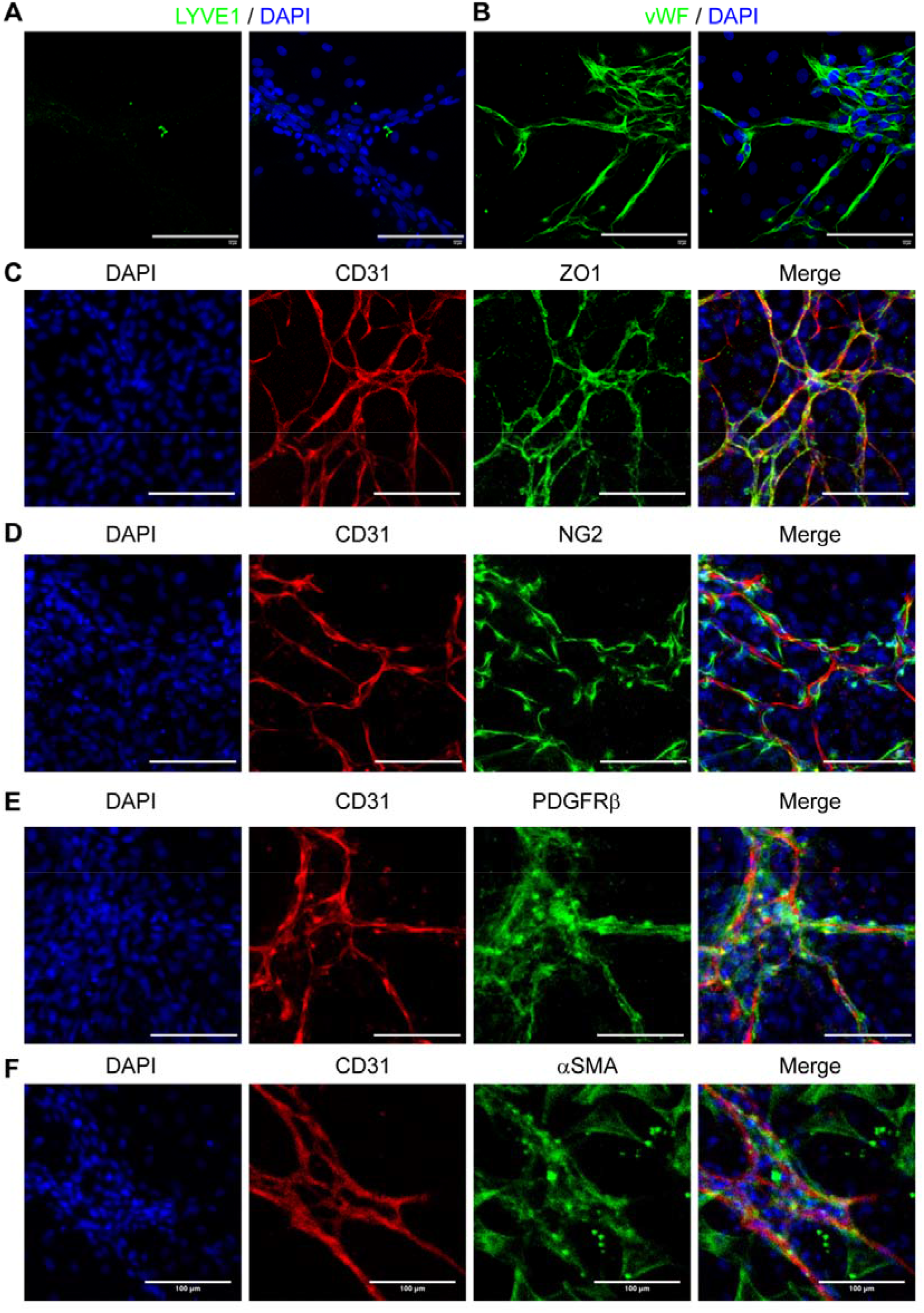
Endothelial and pericyte marker expression in Medium-X. The microvessels were immunostained by the antibodies against LYVE1 (A), vWF (B), CD31 (C-F), ZO1 (C), NG2 (D), PDGFRβ (E), and αSMA (F). Nuclei were stained by DAPI. Scale bars, 100 μm.

TGFβ signaling pathway is essential for the differentiation and maturation of vascular mural cells, including smooth muscle cells and pericytes^38^. There were still some NG2+ pericytes around microvessels in a week of culture in Medium-X containing A83-01 (Fig. 5). Thus, we investigated whether NG2+ pericytes still existed in a long-term culture of 2 weeks, when TGFβ signaling was inhibited by different concentrations of A83-01. We found that higher concentrations of A83-01 induced a higher density of microvessels and the concentration of 0.5 μM gave the maximal level of microvessel length and branch formation (Fig. 6), which was consistent with previous report^39^. However, there were few NG2+ pericytes when A83-01 concentration was bigger than 0.2 μM. (Fig. 6). Restoration of the TGFβ signaling by removing A83-01 from the medium at the second week rescued NG2+ pericytes to some extent (Fig. 7). However, NG2+ pericytes cannot be recovered if A83-01 was removed at the third week (data not shown).

**Fig 6.**
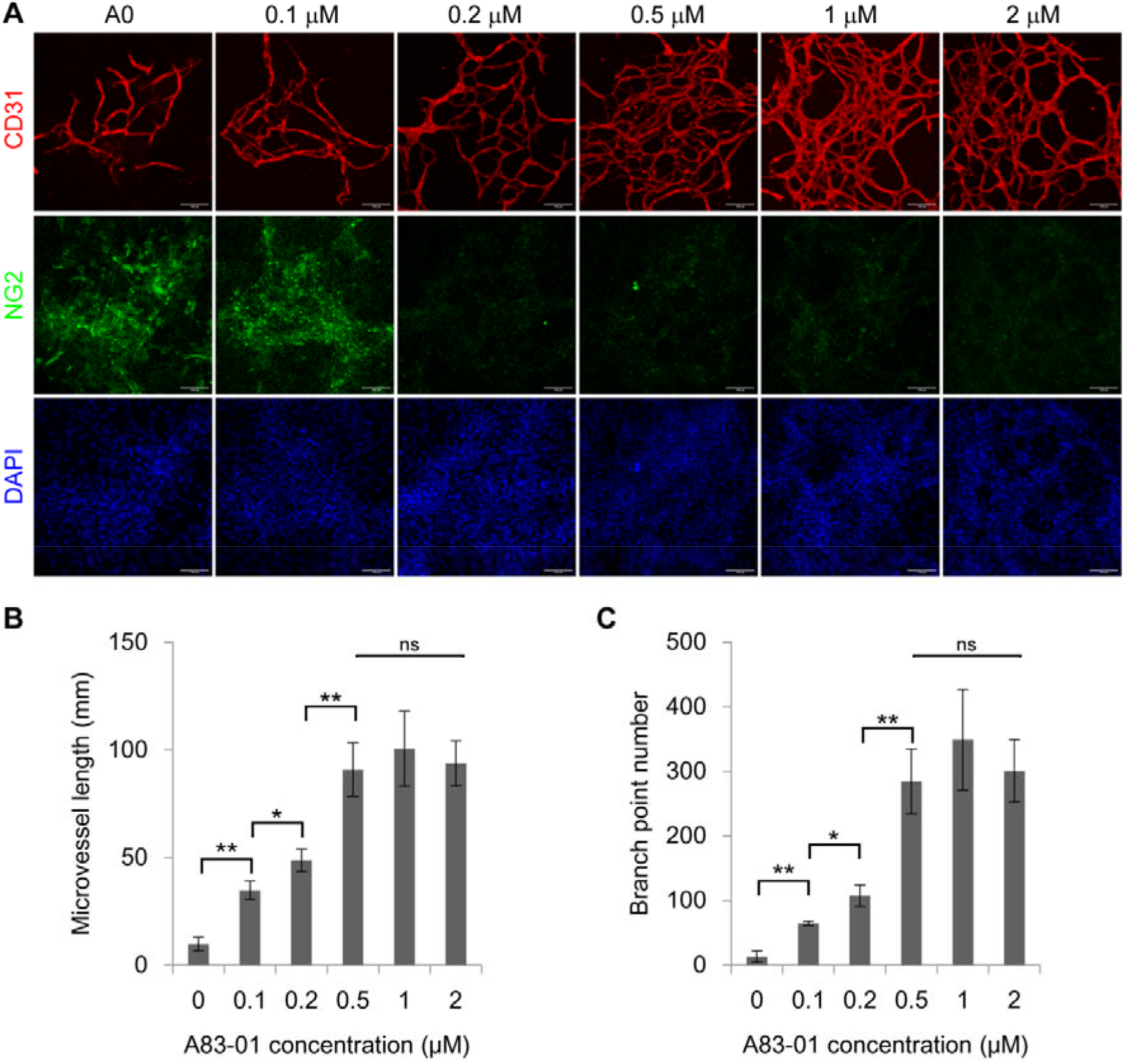
The effect of TGFβ inhibitor A83-01 on microvessels. A, The primary microvessels were cultured in the media supplemented with different concentrations of A83-01, including 0, 0.1, 0.2, 0.5, 1, 2 μM, for 2 weeks, and immunostained by the antibodies against CD31 and NG2. Nuclei were stained by DAPI. Scale bars, 100 μm. B and C, Quantification of microvessel length and branch point number in different groups. Data were presented as mean ± SD. One-way ANOVA was performed on the data, followed by Bonferroni post hoc tests. *p < 0.05, **p < 0.01.

**Fig. 7.**
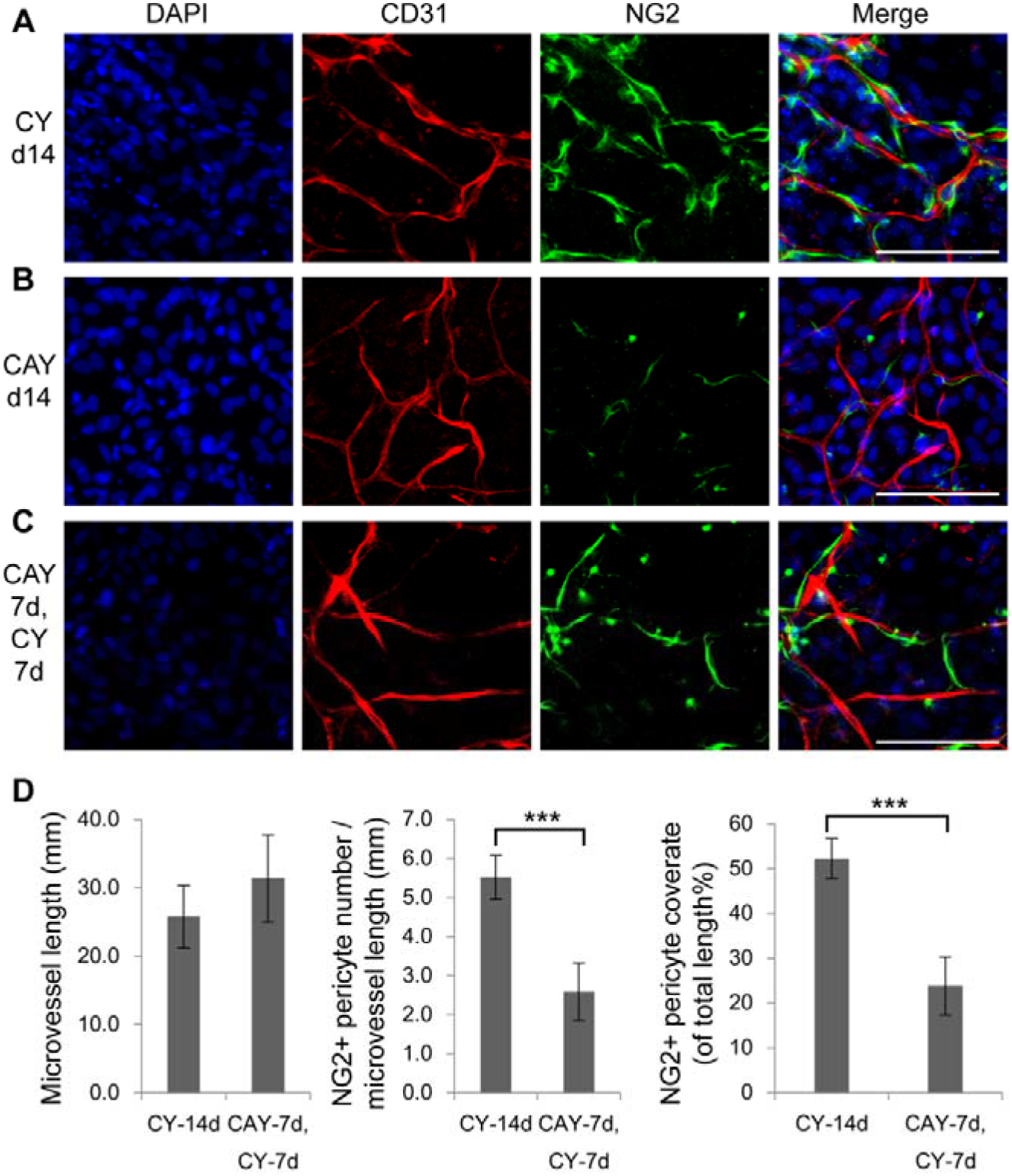
Dynamic regulation of NG2 pericytes by A83-01. A-C, The primary microvessels were cultured in CY for 14 days (A), in CAY for 14 days (B), and in CAY for 7 days, followed by CY for another 7 days (C), then immunostained by antibodies against CD31 and NG2. Nuclei were stained by DAPI. Scale bars, 100 μm. D, Quantification of microvessel length, NG2+ pericyte number per microvessel length, and NG2+ pericyte coverage ratio of total microvessel length (%). Data were presented as mean ± SD. Student’s *t*-test was performed to analyze significant differences between groups. ***, p < 0.001.

## 3. Discussion

This study presented a model platform for expanding tubular microvessels with syngeneic ECs and pericytes on the rigid surface of regular culture dishes. The subcutaneous soft connective tissue of the adult body contains microvessels, fibroblasts, and some stem cells^40,41^. It can be relatively easily obtained during minor surgeries, such as liposuction, plastic surgeries, and foreskin surgeries, making it a promising source for translational medicine. In traditional EC isolation methods, the tissues were digested for a long time until single ECs were obtained. Long-term exposure to digestive enzymes may harm the cells to some extent. The isolated single ECs lost their microenvironment, including cell-cell connections between ECs and between ECs and pericytes. In this study, we developed a cyclic digestion method to maximize the viability of primary cells and the structural integrity of microvessels. This method can be applied to primary cell isolation from other tissues and organs in general.

The maintenance of tubular morphology of adult microvessels in this study was achieved through the orchestration of multiple pathways. One of the main threats to the cells cultured *in vitro* is oxidative stress, which induces DNA damage, senescence, and apoptosis^42^. Oxidation was also a major biological process in the culture of EGM2MV (Fig. 2, A). The addition of antioxidants may contribute to the downregulation of oxidation pathways in Medium-X. The GSK3β inhibitor Chir99021 upregulated Wnt signaling and played a dual role in the system. Activated Wnt signaling has been reported to suppress adipogenesis^12^. On the other hand, Wnt signaling was a master regulator in angiogenesis, blood vessel morphogenesis during embryonic development and adult tissue regeneration^43^. Our results showed that the treatment of Chir99021 alone could activate most of the proangiogenic genes of adult microvessels (Fig. 3, C and E), indicating that Wnt signaling also played a central role in the tubular microvessel growth *in vitro*.

TGFβ signaling pathway is essential for vascular development as it modulates the differentiation and maturation of vascular mural cells, including smooth muscle cells^44^ and pericytes^45^. On the other hand, TGFβ signaling can induce the loss of EC identity through endothelial-to-mesenchymal transition (EndMT)^13^. TGFβ inhibition maintained EC identity and proliferation^39^. Our results showed that the inhibition of TGFβ signaling by A83-01 upregulated the expression CD34, a marker for endothelial progenitor cells^46^. Treatment of A83-01 also promoted the expression of the genes for EC proliferation, such as Pgf, Igf1, and Adm. More importantly, the critical functional genes, such as Apoe for cholesterol transport and Thbd for anti-coagulation, were upregulated by the treatment of A83-01 (Fig. 3). All these results indicate that the inhibition of TGFβ signaling by A83-01 promoted EC proliferation and maintained EC functions.

Y27632 is a ROCK inhibitor that has been widely used in the culture of pluripotent stem cells and functions to promote cell survival *in vitro*^14^. Studies in corneal endothelium showed that Y27632 promoted cell proliferation by upregulating Cyclin D and downregulating p27 gene expression^47^. However, few genes were influenced by the addition of Y27632 in our study (data not shown), indicating that Y27632 may take effect at the protein level. Y27632 has been reported to reduce cell contractility by down-regulating the phosphorylation of non-muscle myosin light chain II (MLC)^48^. As pericytes wrap around microvessels and exert contractile forces to the EC tubules and limit their growth^49^, Y27632 may promote microvascular tubule growth and branch formation by reducing the contractility of ECs and pericytes, which needs further investigation.

There have been several models for engineering tubular microvessels *in vitro*. A classic method is to culture ECs on the surface of Matrigel, and ECs will self-assemble into tubules and form a network, which, however, can only survive a few days^3^. The ECs can also self-assemble into tubular structures inside a 3D matrix, such as collagen, Matrigel, and fibrin gels^3^. Microfluidics techniques can make these 3D microvessels perfused, better-mimicking microvessels *in vivo*^3^. These tubular microvessels in 3D hydrogels were usually maintained and investigated for up to 2 weeks^3^.

Most microvascular diseases are chronic diseases and take place in years or decades. The potential effect of a treatment may not emerge in a short time. A goal in the research of microvascular engineering is to maintain the culture of microvessels for a longer time. In this study, the microvessels in the experimental groups of C, CA, CY, and CAY had minor differences at the first week time point, but they behaved distinctly from the second to eighth week (Fig. 4). The microvessels in group C quickly deteriorated in the second week, indicating that the activated Wnt signaling pathway alone promoted angiogenesis but was not enough to maintain the structural stability of microvessels *in vitro*. The beneficial effect of Y started to emerge from the second week. Adding Y to the media C or CA significantly improved microvessel length and branch formation. The addition of A to CY medium further promoted microvessel stability, possibly through upregulating the genes related to EC functions and proliferation (Fig. 3). The microvessel length and branch formation in Medium-X (VEGF+CAY) were maintained at a relatively high level for at least 4 weeks *in vitro* (Fig. 4). The time period of the third and fourth weeks had relative stable microvessels, which are a proper candidate for experimental manipulations (Fig. 4).

There was a subtle balance between the ECs and pericytes in this model. Pericytes play a vital role in regulating microvessel structural stability and functions^36^. NG2 is a glycoprotein located on the plasma membrane of pericytes and regulates microvessel integrity^36^. Genetic knock-out studies showed that NG2^-/-^ animals had fewer microvessels than normal controls^50^. Our results are consistent with previous reports that TGFβ signaling is essential for the differentiation and maturation of pericytes^38^. Although there were slight differences in NG2+ pericytes between CY and CAY media at the first week, a long-term culture of 2 weeks in CAY medium significantly reduced the number of NG2+ pericytes around microvessels (Fig. 6 and 7). NG2+ pericytes can be preserved by removing A83-01 from the medium from the second week (Fig. 7), but they will vanish if A83-01 was removed from the third week. These results indicate that the maintenance of NG2+ pericytes requires TGFβ signaling. On the other hand, TGFβ signaling played an opposite role in ECs. TGFβ inhibition was required to for maintaining EC identity and functions *in vitro*^13,39^. Future studies are warranted to investigate how to coordinate the growth of syngeneic ECs and pericytes in tubular microvessels *in vitro*.

## 4. Outlook

Most microvascular studies use the coculture models of ECs and pericytes from different tissue sources. Given the heterogeneity of the vascular system, our model will hopefully provide more insights into the mechanisms of EC-pericyte interactions in tissue regeneration and diseases. The method established in this study also applies to other organs for engineering organ-specific microvascular models. As human microvessels can be relatively easily obtained from the subcutaneous soft connective tissues during minor surgeries, the platform presented in this study can hopefully be translated to clinical settings, benefiting the development of personalized microvessel models for precision medicine.

## 5. Materials and methods

### Isolation of primary microvessels from adult rats

The animal procedure was conducted according to the Ministry of Science and Technology guide for laboratory animal care and use and approved by the animal care and use committee of Qingdao University. Male Sprague Dawley (SD) rats aged 8–10 weeks were euthanized by an overdose of isoflurane and bilateral thoracotomy. The subcutaneous soft connective tissue was harvested in an aseptic environment. The tissue pieces were digested in an enzymatic solution containing 2 mg/ml collagenase I (Wathington, Cat#LS004196), 5 mg/ml bovine serum albumin (BSA, Sigma, Cat#A1933), and 5 μM Y27632 (Sellect, Cat#S1049) in DMEM (Invitrogen, Cat#12800017).

The cyclic digestion method was used to harvest primary microvessels. Every 10 minutes, the cell suspension was harvested, a new enzymatic solution was added to the remaining tissue, and the digestion cycle continued until all the tissue pieces were digested. The cell suspension was centrifuged at 1000 rpm for 4 minutes, and cell pellets were re-suspended in phosphate-buffered saline (PBS). All the cell pellets were pooled together and filtered through a 30 μm strainer to remove red blood cells and single cells. The long microvessel segments were collected from the strainer mesh and used for culture.

### Culture of primary microvessels

The primary microvessel segments were cultured in regular plastic dishes and plates in different media. EGM2MV medium was purchased from Lonza (Cat# CC-3202). Our custom-made medium was DMEM/F12 (Invitrogen, Cat# 12400024), supplemented mainly with 2% fetal bovine serum (FBS, Gibco, Cat# 10091148), 100 U/ml penicillin and 100 μg/ml streptomycin (Gibco, Cat#15140122), 5 mM nicotinamide (Sigma, Cat#N0636), 1 mM NAC (Sigma, Cat#A9165), 50 μM Vitamin C (Sigma, Cat#A4403), 3 μM glutathione (Sigma, Cat#G6013), 10 μg/ml insulin (Aladdin, Cat#I113907), 7.5 μg/ml transferrin (Sigma, Cat#T0665), 40 nM Sodium Selenite (Sigma, Cat#S9133).

Medium-X was supplemented with 2 μM Chir99021 (Sellect, Cat#S2924), 0.2 μM A83-01 (Tocris, Cat#2939), 5 μM Y27632 (Sellect, Cat#S1049), and 10 ng/ml VEGF (Peprotech, Cat# 100-20). For the titration of A83-01, the concentrations included 0, 0.1, 0.2, 0.5, 1, and 2 μM.

The cells were cultured in an incubator at 37 °C, with 5% CO_2_ and 95% humidity. The medium was changed every other day in the first four days and every day after that.

### Total RNA extraction

Total RNA was extracted from the microvascular cells using Trizol (Invitrogen, Cat#15596026) according to the manufacture instruction. The cells of a 60 mm dish were homogenized in 1 ml Trizol, centrifuged at 12,000×g for 5 minutes at 4°C, and then the supernatant was transferred into a tube with 0.3 mL chloroform/isoamyl alcohol (24:1). The mixture was centrifuged at 12,000×g for 10 minutes at 4°C, the upper aqueous RNA layer was transferred into a new tube with equal volume of isopropyl alcohol, and then centrifuged at 12,000×g for 20 minutes at 4°C. The RNA pellet was washed with 1 ml 75% ethanol, and then centrifuged at 12,000×g for 3 minutes at 4°C. The pellet was dried in air for 5-10 minutes. Finally, 50 μl DEPC water was added to dissolve the RNA, which was qualified and quantified using a Nano Drop and Agilent 2100 bioanalyzer (ThermoFisher).

### mRNA Library Construction

The mRNA was purified by Oligo(dT)-attached magnetic beads and fragmented into small pieces. First-strand cDNA was generated by random hexamer-primed reverse transcription and a second-strand cDNA synthesis. A-Tailing Mix and RNA Index Adapters were added. The cDNA fragments were amplified by PCR, and then purified by Ampure XP Beads and dissolved in EB solution. The product was validated on an Agilent Tech 2100 bioanalyzer. The double stranded PCR products were denatured and circularized by the splint oligo sequence to make the final library, which was amplified with phi29 to make DNA nanoball with more than 300 copies of one molecular, which were loaded into the patterned nanoarray and pair end 150 bases reads were generated on DNBSEQ-T7 platform.

### Sequencing analysis

The sequencing data was filtered with SOAPnuke (v1.5.2). The clean reads were mapped to the reference genome using HISAT2 (v2.0.4). Bowtie2 (v2.2.5) was applied to align the clean reads to the reference coding gene set, then expression level of gene was calculated by StringTie (v2.1.2). KEGG and GO analysis were performed in DAVID Bioinformatics Resources (https://david.ncifcrf.gov). The heatmap and bubble plot were drawn by SRplot (http://www.bioinformatics.com.cn) according to the gene expression in different samples.

### Quantitative RT-PCR analysis

Total RNA was extracted from the microvascular cells using Trizol (Invitrogen, Cat# 15596026) according to the manufacture instruction. RNA quality was checked by measuring the optical density ratio at 260/280 nm on a nanodrop. RNA from each sample was reverse transcribed into cDNA by an Evo M-MLV RT Mix Kit (Accurate Biotechnology, Cat#AG11728). Experiment was carried out using the forward and reverse primers listed below. Fold change in mRNA expression levels was calculated by the comparative Ct method, using the formula 2-(ΔΔCt) and GAPDH as a calibrator. The primer list:

Apln (AAGCCCAGAACTTCGAGGAC;GGCAGCATATTTCCGCTTCTG),
Dll4 (AGTGTACTCCCGCACTAGCC;CGATGCCTCGGTAGGTAATCC),
Kcne3 (ACAGATCGCAGAGTCAGATCAC;TGATTGTCTGGCCCTGTTCC),
Pcdh12 (CAGCAGGTCTGAAGTGGGAG;GTAGCATCGTGCTTACCGGA),
Plxnd1 (ATCGCCCAGGCCTTCATAGAT;TCTTCCGGTACTCGGGGAT),
Car2, (GCTGGAATGTGTGACCTGGA;CCCAGCTGCAGGGTCATTTT),
Wnt5a (TGGGCACATTTCCACGCTAT;TGTCCTTGAGAAAGTCCCGC),
Fzd1 (GCCTCACAACCAGTCCACAA;TGCTTTACAAATGCCACTCGG),
Cxcl12 (AGCCTTAAACAAGAGGCTCAAG;TGAGGGTGGATCTCGCTCTT),
Mmp9 (GGATCCCCCAACCTTTACCAG;AAGGTCAGAACCGACCCTACA).

### Immunostaining

The cells were fixed in 4% paraformaldehyde (PFA) for 30 min, washed with PBS for three times, permeabilized by 0.1% Triton for 10 minutes, blocked in 5% normal donkey serum for an hour, and incubated with primary antibodies in the blocking solution at room temperature. After 2 hours of primary antibody incubation, the cells were washed in PBS for three times and incubated with secondary antibodies in blocking solution for 1 hour at room temperature. The primary antibodies used in this study included CD31 (Abcam, Cat#ab28364; Santa Cruz, Cat#SC376764), vWF (Proteintech, Cat#11778-1-AP), LYVE1 (Affinity, Cat#AF4202), NG2 (Proteintech, Cat#55027-1-AP), ZO1 (Proteintech, Cat#21773-1-AP), PDGFRβ (Proteintech, Cat#13449-1-AP), and SMA (Santa Cruz, Cat#sc-32251). The secondary antibodies used in this study were purchased from Invitrogen (Cat#A10040, A-21202). Cell nuclei were stained by DAPI (4’,6-diamidino-2-phenylindole). Confocal imaging was performed on a Leica SP8 confocal microscope.

### Statistical analysis

To quantify the features of the microvessels, at least 3 images randomly selected from each group were analyzed using ImageJ. The total microvessel length and branch point numbers used for analysis were the sums on a 10× image area. Data were presented as means ± SD unless otherwise indicated. Student’s *t*-test was used to analyze the differences between groups.

## Credit author statement

DW designed the experiments. DW and JL wrote the manuscript. XS, YY, YL, YX, and DW performed primary cell isolation, cell culture, immunostaining and confocal microscopy. JM performed qRT-PCR. ZW performed quantification of microvessels. HZ, XQ, PL, JL, and DW performed transcriptomic analysis.

## Data availability statement

The raw/processed data required to reproduce these findings cannot be shared at this time due to technical or time limitations. They can be made available upon request.

## Acknowledgements

This work was funded by the Natural Science Foundation of Shandong Province (No. ZR2019LZL001), National Nature Science Foundation of China (No. 32101020, 91849209, and 8187020984), and People’s Livelihood Science and Technology Project of Qingdao (No. 20-3-4-41-nsh). This work was also supported by the Key Laboratory of Nucleic Acid Biology in Cardiovascular Disease, Shandong Province, and the International Research Centre for Non-coding RNAs and Translational Medicine, Qingdao. Thanks to Zhishang Chang, Qian Wen, and Xuxia Song (the Laboratory of Biomedical Center, Qingdao University) for their technical help in confocal microscopy.

