## Supplemental figures for "Expanding adult tubular microvessels on stiff substrates with endothelial cells and pericytes from the same tissue"

Fig. S1. Quantification of primary microvessels.

Fig. S2. Transcriptomic analysis of gene expression profiles in different groups.

Fig. S3. GO analysis (BP, CC and MF) of the genes upregulated by Chir99021 and A83-01.

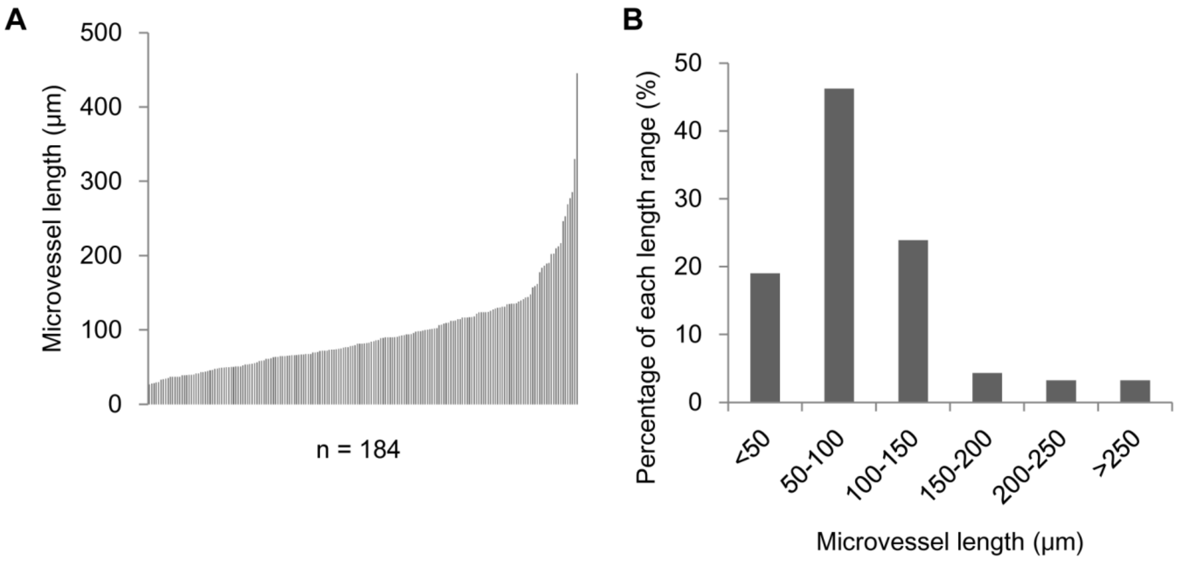


Fig. S1. Quantification of primary microvessels. A, The measurement of fresh primary microvessels isolated from rat subcutaneous soft connective tissues (n=184). B, Percentage of each microvessel length range.

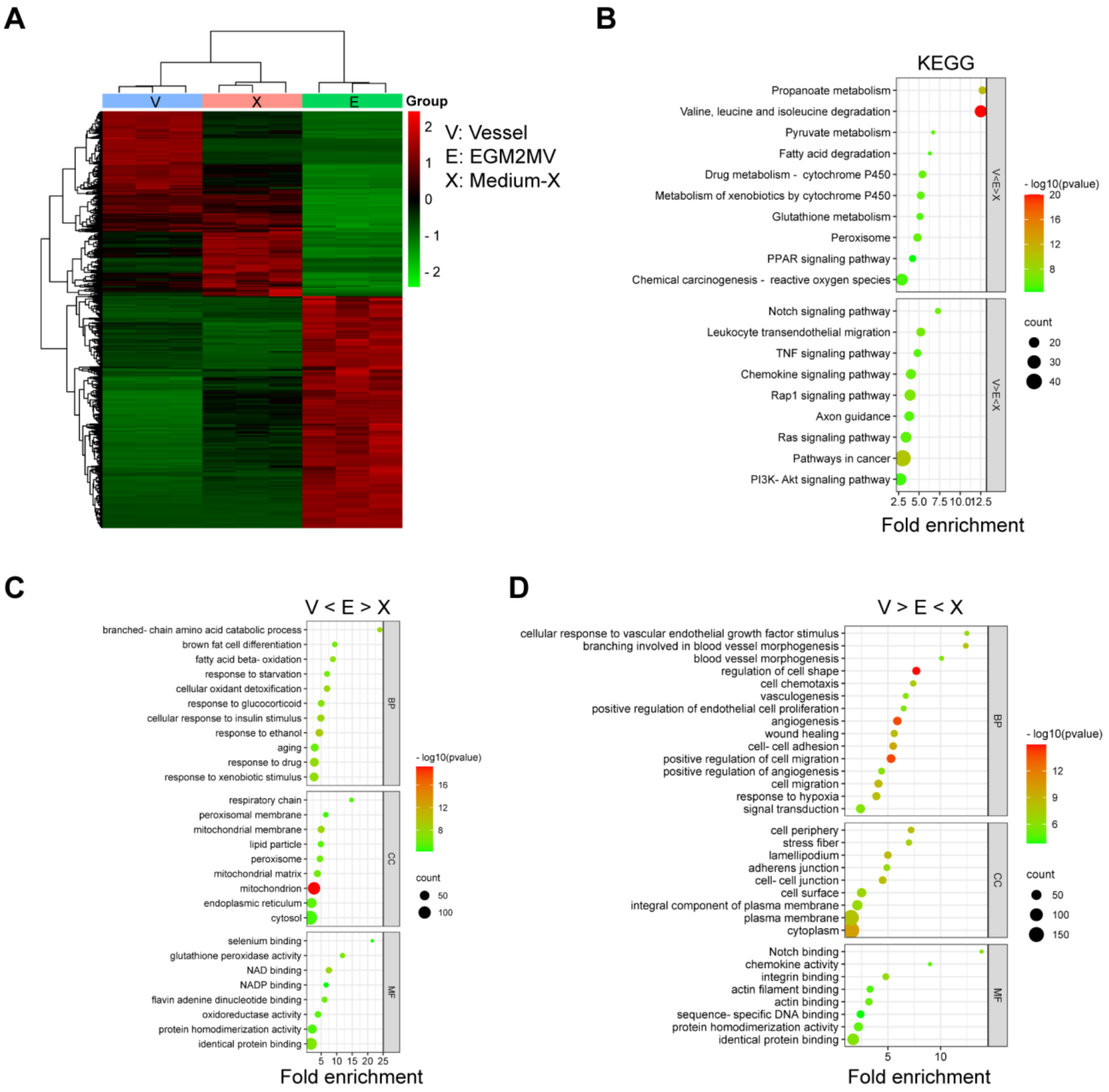


Fig. S2. Transcriptomic analysis of gene expression profiles in different groups. A, Heatmap of the genes up- or down-regulated in EGM2MV and Medium-X. B, KEGG analysis of signaling pathways enriched in EGM2MV and Medium-X. C and D, GO analysis (BP, CC and MF) of the genes enriched in EGM2MV (C) and Medium-X (D).

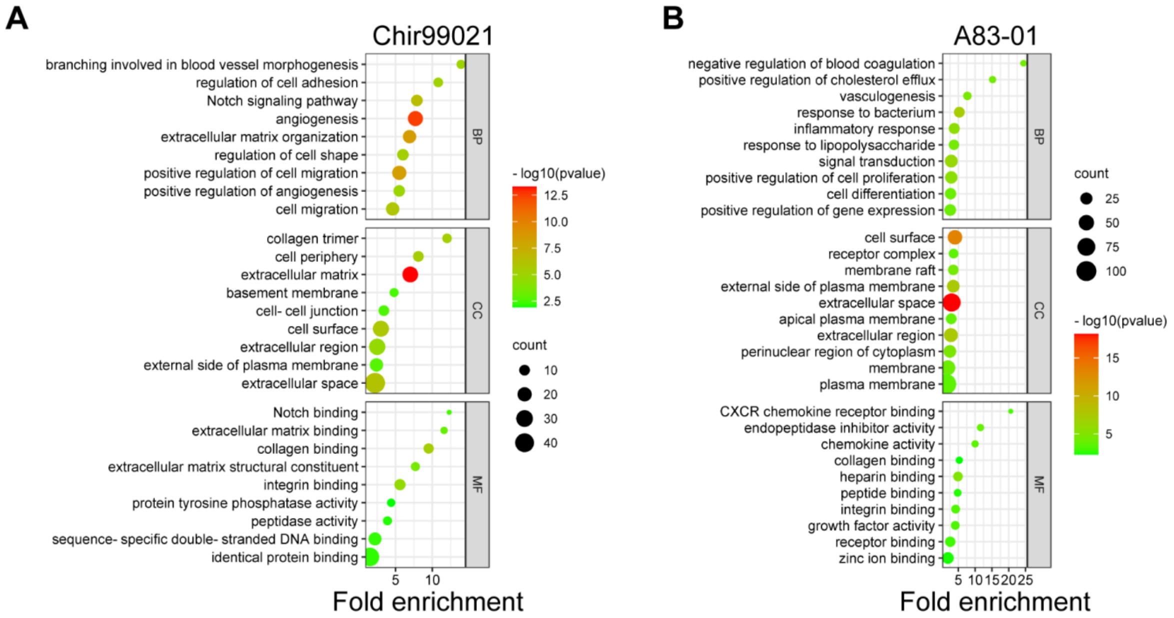


Fig. S3. GO analysis (BP, CC and MF) of the genes upregulated by Chir99021 (A) and A83-01 (B).
